# Biased gene conversion drives codon usage in human and precludes selection on translation efficiency

**DOI:** 10.1101/086447

**Authors:** Fanny Pouyet, Dominique Mouchiroud, Laurent Duret, Marie Sémon

**Affiliations:** Laboratoire de Biométrie et Biologie Evolutive, Université de Lyon, Université Claude Bernard, CNRS UMR 5558, 43 boulevard du 11 novembre 1918, F-69100, Villeurbanne, France; Laboratory of Biology and Modelling of the Cell, Université de Lyon, Université Claude Bernard, CNRS UMR 5239, INSERM U1210, 46 allée d’ltalie, F-69007, Lyon, France

**Keywords:** Codon Usage, Biased gene conversion, Translational selection, Recombination, Meiosis

## Abstract

In humans, as in other mammals, synonymous codon usage (SCU) varies widely among genes. In particular, genes involved in cell differentiation or in proliferation display a distinct codon usage, suggesting that SCU is adaptively constrained to optimize translation efficiency in distinct cellular states. However, in mammals, SCU is known to correlate with large-scale fluctuations of GC-content along chromosomes, caused by meiotic recombination, via the non-adaptive process of GC-biased gene conversion (gBGC). To disentangle and to quantify the different factors driving SCU in humans, we analyzed the relationships between functional categories, base composition, recombination, and gene expression. We first demonstrate that SCU is predominantly driven by large-scale variation in GC-content and is not linked to constraints on tRNA abundance, which excludes an effect of translational selection. In agreement with the gBGC model, we show that differences in SCU among functional categories are explained by variation in intragenic recombination rate, which, in turn, is strongly negatively correlated to gene expression levels during meiosis. Our results indicate that variation in SCU among functional categories (including variation associated to differentiation or proliferation) result from differences in levels of meiotic transcription, which interferes with the formation of crossovers and thereby affects gBGC intensity within genes. Overall, the gBGC model explains 70% of the variance in SCU among genes. We argue that the strong heterogeneity of SCU induced by gBGC in mammalian genomes precludes any optimization of the tRNA pool to the demand in codon usage.

## Introduction

Although synonymous codons encode the same amino acid, some are used more frequently than others. This preferential usage of a subset of synonymous codons is known as “codon usage bias”. In many species, including humans, this bias varies substantially among genes in the genome. Both adaptive and non-adaptive processes, which are not mutually exclusive, have been proposed to explain the existence of codon usage biases (Chamary, Parmley, & Hurst, 2006; Duret, 2002; Plotkin & Kudla, 2011). According to the main adaptive model, termed translational selection, synonymous codon usage (SCU) and abundance of tRNA are co-adapted to optimize the efficiency of translation (Dos Reis & Wernisch, 2009; Drummond & Wilke, 2008; Hershberg & Petrov, 2008; Ikemura, 1981; Kanaya, Yamada, Kinouchi, Kudo, & Ikemura, 2001). Non-adaptive models propose instead that codon usage bias results from biases in neutral substitution patterns, driven by mutation or by GC-biased gene conversion – gBGC, (Chen, Lee, Hottes, Shapiro, & McAdams, 2004; Duret & Galtier, 2009; Galtier, Piganeau, Mouchiroud, & Duret, 2001; Sémon, Lobry, & Duret, 2006). In humans, these processes have been long studied, but the relative influence of adaptive and nonadaptive processes on SCU is still a matter of debate (Chamary et al., 2006; Duret, 2002; Plotkin & Kudla, 2011).

Recently, Gingold et al., (2014) compared synonymous codon usage among sets of human genes associated to different functional categories (as defined in the Gene Ontology) and observed a striking difference between sets of genes associated with cellular proliferation and those associated with differentiation. They also observed that the relative abundance of tRNA varies according to the proliferative or differentiative state of cells, which was logically interpreted in term of translational selection: different cell types express specific sets of genes whose coding sequence is co-adapted with specific pools of tRNAs (Gingold et al. 2014).

However, this interpretation stands in contradiction with two other studies. First, expression levels of individual tRNA genes do indeed vary extensively between tissue types and developmental stages in mice. But when tRNA genes are grouped by isoacceptor families (which recognize the same codon) the resulting collective expression levels are stable throughout development and specify a constant pool of anticodons (Schmitt et al. 2014). Second, in continuation to this work, a recent study specifically contrasted cells undergoing proliferation and those undergoing differentiation, and found no covariation of tRNA pool and codon usage between these cells (Rudolph et al. 2016). Hence, neither result is consistent with the initial claim interpreting differences in codon usage bias between functional classes as a consequence of translational selection. Then, why does synonymous codon usage vary between genes associated to different functional categories?

It has long been known that in mammals, variation in synonymous codon usage between genes is linked to large-scale fluctuation of GC-content along chromosomes, the so-called isochores (Bernardi et al. 1985; Mouchiroud et al. 1988; Mouchiroud et al. 1991; Clay and Bernardi 2011). There is strong evidence that isochores are the consequence of GC-biased gene conversion (gBGC), a form of segregation distortion that occurs during meiotic recombination and that favors the transmission of GC alleles over AT alleles (Duret & Galtier, 2009; Munch, Mailund, Dutheil, & Schierup, 2014; Williams et al., 2015). The gBGC process leads to an increase in the GC-content in regions of high recombination rate, which affects both coding and non-coding regions, including synonymous codon positions (Duret & Galtier, 2009; Galtier & Duret, 2007; Glémin et al., 2015). Besides, it has been shown that the rate of recombination within genes is negatively affected by their level of expression in the germline (McVicker and Green 2010). This suggests that the impact of gBGC on the synonymous codon usage of genes might depend on their pattern of expression.

To investigate the parameters responsible for variation in codon usage among functional categories, we analyzed the relationships between synonymous codon usage, GC-content, recombination rate and expression patterns of human genes. We first show that the variation in codon usage among functional categories results from differences in GC content. Then we propose a new test to show that it is not associated with translational selection. Instead, synonymous codon usage correlates with large-scale variation in genomic GC-content and with differences in intragenic recombination rate. In turn, the difference in intragenic recombination rate between functional categories is explained by their expression level in meiosis. Altogether, GC-content of non-coding regions and meiotic expression explain 70% of the variation in synonymous codon usage of human genes.

At the end, our results are fully consistent with the hypothesis that synonymous codon usage is driven by gBGC, and not by translational selection, and that the differences observed among functional categories reflect variation in long-term intragenic recombination rates, resulting from differences in meiotic expression levels.

## Results

### Variation in codon usage among functional categories results from differences in GC-content

To better understand what distinguishes codon usage between sets of genes involved in cellular proliferation and differentiation, we started by investigating the main factors that would discriminate codon usage between functional categories in general. For this purpose, we grouped genes per functional category (687 biological processes, associated to more than 40 genes in the Gene Ontology database), and computed codon frequencies for each of these gene sets. Variation in codon usage among these GO gene sets was analyzed by Principal Component Analysis (PCA). In agreement with Gingold *et al.* (2014), the first principal component of this analysis explains 41% of the total variance, and segregates “proliferation” (7 categories) from “differentiation” (6 categories) GO categories (Fig S1A). To remove potential confounding effects of variation in amino acid composition between these categories, we also analyzed the relative synonymous codon usage (RSCU; see material and methods) for each functional category. The first principal component of the PCA analysis is even stronger, explaining 73% of the total variance of RSCU, and still separates categories related to proliferation (red dots) and differentiation (blue dots, Fig 1A). Thus, synonymous codon usage clearly varies between functional categories in general, and between proliferation and differentiation in particular. What property of sequence composition underlies this difference? It is well known that synonymous codon usage is strongly correlated to GC content at third position of codons – termed GC3; (Mouchiroud et al. 1988). Thus, we computed the average GC3 of each GO gene set. The GC3 of these functional categories vary widely (from 0.45 to 0.73) and is perfectly correlated to their coordinates on the first PCA axis (R^2^ = 0.99; Fig 1B). Hence, variation in SCU between functional categories is fully explained by variation in GC3.

**Fig 1:**
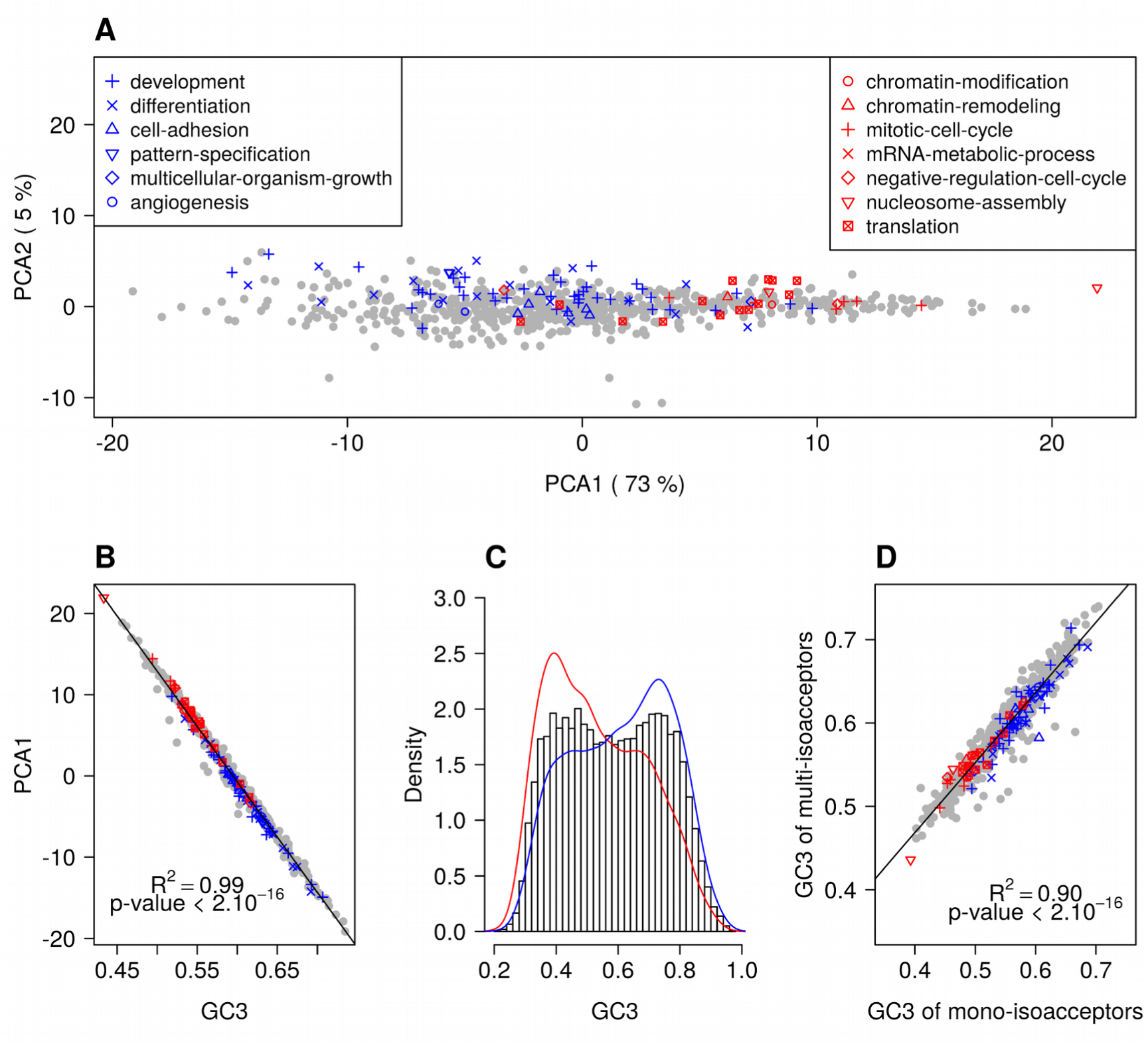
Variation in synonymous codon usage and in GC3 among functional categories. (A) Factorial map of the principal-component analysis of synonymous codon usage in GO functional categories in the human genome. Each dot corresponds to a GO gene set, for which the relative synonymous codon usage (RSCU) was computed. GO categories that are associated with “differentiation” or with “proliferation” are displayed respectively in blue and in red. (B) Correlation between the RSCU of GO gene sets (first PCA axis) and their average GC-content at third codon position (GC3). (C) Distribution of GC3 of human protein coding genes. The black histogram represents the distribution for whole dataset (15,970 genes). The blue curve (resp. the red curve) is a smoothed distribution of GC3 for “differentiation” genes (N=2,833) (resp. “Proliferation” genes, N=1,008) (D) Correlation between the GC3 of mono-isoacceptor amino-acids and multi-isoacceptor amino-acids. For each GO gene set, the average GC3 was computed separately for amino-acids decoded by multiple tRNA isoacceptors (N=14 multi-isoacceptor amino-acids), and for those decoded by one single tRNA isoacceptor (mono-isoacceptor amino-acids: Phe, Asp, His, Cys). Amino-acids encoded by a single codon (Met, Trp) were excluded.

On average, in our dataset, each gene is associated to 9 GO biological processes. Many genes belong to more than one GO biological-process category, either because they have several functions (pleiotropy) or because these categories are nested from specific to broad functions. Hence, GO-terms are not independent. To avoid this redundancy, for the remainder of this study we switched from analyses at the level of GO gene sets to analyses at the level of individual genes (except when stated otherwise). Each gene was assigned with one of three categories based on their GO annotation: 1,008 genes associated with “proliferation”, 2,833 genes associated with “differentiation”, and 12,129 “other” genes unrelated to these keywords (see methods). The distribution of GC3 content over the entire dataset is bimodal (Fig 1C). For the subsets of genes associated to “proliferation” and “differentiation”, the two distributions of GC3 differ significantly from each other (T-test, p-value < *2.10^−16^*), and their peaks coincide with each of the two modes observed for the whole genome. Genes associated to “proliferation” are on average less GC-rich than genes associated to “differentiation” (mean GC3 are respectively 0.53 and 0.61 in the two subsets).

### Variation in synonymous codon usage is not driven by translational selection

We first investigated whether the observed variation in synonymous codon usage (i.e. variation in GC3) might be driven by translational selection. This model proposes that the relative usage of synonymous codons should co-vary with the abundance of their cognate tRNAs. A property of the tRNA gene repertoires allows us to test this hypothesis. The human genome contains 506 tRNA genes (decoding the 20 standard amino-acids), corresponding to 48 different tRNA isoacceptors (Chan and Lowe 2009). Among the 18 amino acids having two or more synonymous codons, four are decoded by a single tRNA isoacceptor (mono-isoacceptor amino-acids: Phe, Asp, His and Cys), and the 14 other ones are decoded by several tRNA isoacceptors (multi-isoacceptors amino-acids).

For multi-isoacceptors amino-acids, the relative abundance of the different tRNA isoacceptors can vary among different cell types, and hence might covary with the relative synonymous codon usage of genes preferentially expressed in these cell types. For instance, let us consider Gln, which has two synonymous codons (CAG, CAA) that are decoded by two tRNA isoacceptors (respectively anticodons CTG and TTG). Let us consider a theoretical example of two cell types (say A and B) that differ in their relative tRNA abundance (CTG-tRNA being more abundant in A cells, and TTG-tRNA in B cells). According to the translational selection model, sets of genes that are over-expressed in A cells, should preferentially use the CAG codon whereas genes that are over-expressed in B cells, should preferentially use the CAA codon. However mono-isoacceptor amino-acids are, by definition, decoded by a single tRNA isoacceptor and the relative tRNA abundance cannot vary across cell types. Hence, according to the translational selection model, the relative synonymous codon usage for mono-isoacceptor amino-acids is not expected to vary among cell-specific gene sets. In other words, for mono-isoacceptor amino-acids, variation in synonymous codon usage among GO gene sets cannot be explained by co-adaptation with the tRNA pool.

To test whether variation in synonymous codon usage was driven by translational selection, we computed synonymous codon usage (GC3) in GO gene sets, separately for codons corresponding to mono-isoacceptor amino-acids and for codons corresponding to multiisoacceptor amino-acids. We observed that the range of variation in GC3 is very similar for mono- and multi-isoacceptor amino-acids. Importantly, the two parameters are strongly correlated (*R^2^* = 0.90) (Fig 1D). This implies that GC3 variation is driven by a process that affects both mono-isoacceptor and multi-isoacceptor amino-acids, and hence that this process is not related to variation in tRNA abundance. This observation holds true for all functional categories, including those associated to differentiation or proliferation (red and blue dots in Fig 1D).

### Impact of large-scale variation in genomic GC-content on synonymous codon usage

So far, we have shown that genome-wide variation in synonymous codon usage is not driven by translational selection. Early studies, more than 30 years ago, have shown that variation in human synonymous codon usage is strongly correlated with large-scale fluctuations of GC-content along chromosomes (the so-called isochores), affecting both coding and noncoding regions (Bernardi et al. 1985; Mouchiroud et al. 1988; Mouchiroud et al. 1991; Clay and Bernardi 2011). We therefore tested whether genes associated with “proliferation” were located in genomic regions with a lower GC-content than genes associated with “differentiation”. We observed as expected that the GC3 of genes correlates with the GC-content of their flanking regions (Fig 2A, *R^2^* = 0.48, p-value <2.10*^−16^*). This correlation is observed for all genes, including the subsets of genes associated with “proliferation” and “differentiation” (*R^2^* = 0.48 and 0.46, all p-values <2.10*^−16^*). Thus, in agreement with the literature (Bernardi et al., 1985; Clay & Bernardi, 2011; Mouchiroud et al., 1991; Mouchiroud et al., 1988), our observations indicate that variation in SCU between genes is to a large extent attributable to their position in the genome. However, when the regional GC-content is controlled for, there remains a significant difference in GC3 between gene categories: on average, for a given regional GC-content, there is a gap of 5% to 7% of GC3 between the categories “differentiation” and “proliferation” (Fig 2A, p-value <2.10*^−16^*). This implies that the difference in synonymous codon usage between these gene categories does not result from a preferential location in different isochores.

**Fig 2:**
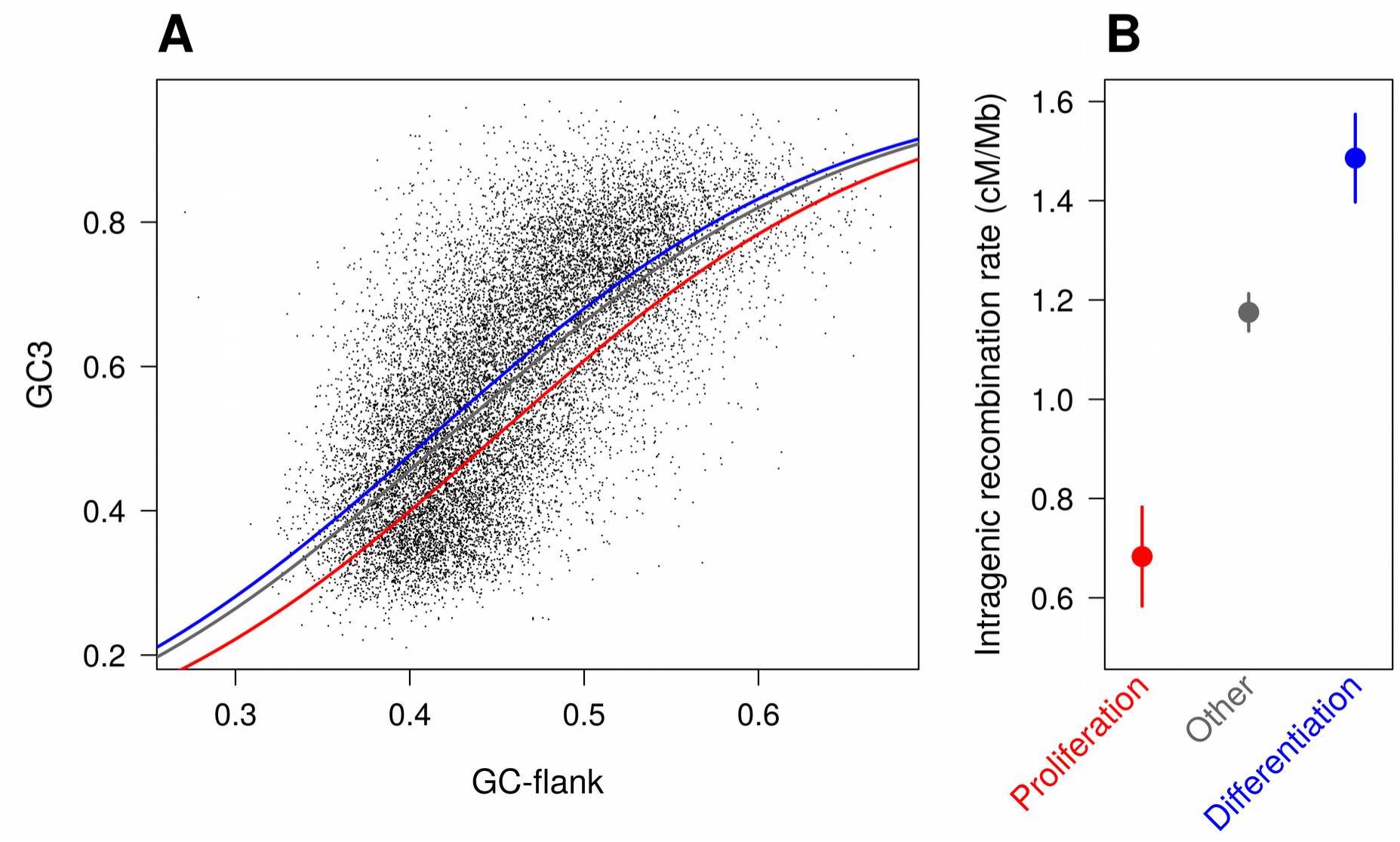
Difference in SCU between “proliferation” and “differentiation” genes is linked to variation in intragenic recombination rate, and not to their isochore context. (A) Correlation between the GC3 of genes and the GC content of their flanking regions (GC-flank). Each dot corresponds to one gene. GC-flank was measured in 10 kb upstream and 10 kb downstream of the transcription unit. The curves show a generalized linear model (glm), predicting GC3 with GC-flank and the gene function. The curves corresponding to “differentiation” genes (blue) “proliferation” genes (red) and other genes (grey) differ significantly (LRT of glm with and without gene function, p-values < 2.10*^−16^*). Correlation coefficients were computed on logit transformed values, independently for “differentiation” genes (N=2,833, R^2^ = 0.46), “proliferation” genes (N=1,008, R^2^ = 0.48), other genes (N=12,129, R^2^ = 0.49) and all genes (N=15,970, R^2^ = 0.48). All p-values < 2.10*^−16^*. (B) Average intragenic recombination rate in each functional category. Error bars represent standard errors.

### Variation in synonymous codon usage among functional categories correlates with differences in intragenic recombination rate

Previous studies have shown that the evolution of GC-content along chromosomes is driven by meiotic recombination, both on a broad (Mb) scale (Duret & Arndt, 2008; Munch et al., 2014) and on a fine (kb) scale (Clément and Arndt 2013; Pratto et al. 2014). There is now strong evidence that this correlation between GC-content and recombination is caused by the process of GC-biased gene conversion (gBGC) which leads to increase the GC-content in regions of high recombination (Duret & Galtier, 2009; Galtier & Duret, 2007;Galtier et al., 2001; Glémin et al., 2015; Munch et al., 2014; Pratto et al., 2014; Williams et al., 2015). Recombination rate varies along chromosomes, and notably tends to be lower within genes than in flanking regions (Myers et al. 2005; McVicker and Green 2010). Interestingly, we observed that intragenic recombination rates (in cM/Mb) differ among the three sets of genes defined previously, and covary with their GC3: the average intragenic recombination rate is lower in “proliferation” genes compared to other genes, whereas it is higher in “differentiation” genes (Fig 2B; p-value of Kruskal-Wallis test < 2.10^*-16*^ as for all pairwise Wilcoxon tests). These observations are therefore consistent with the hypothesis that differences in GC3 between “differentiation” and “proliferation” genes could also be driven by gBGC.

### The difference in intragenic recombination rate between functional categories is explained by their expression level in meiosis

Why do recombination rates vary across functional categories? Previous studies have shown that intragenic recombination rates vary according to gene expression patterns: genes that are expressed in many tissues tend to have lower intragenic crossover rates (McVicker & Green, 2010; Necsulea, Sémon, Duret, & Hurst, 2009). Mc Vicker and Green (2010) analyzed expression levels in many different samples, including both somatic tissues and meiotic or non-meiotic germ cells. Interestingly, they showed that the negative correlation between intragenic recombination rate and expression level is stronger in germ cells than in somatic tissue, and more specifically, stronger in meiotic cells than in other germ cells, most probably because gene expression in meiotic cells interferes with the formation of crossovers (McVicker and Green 2010).

To test whether the differences in intragenic recombination rates that we observed between “proliferation” and “differentiation” genes could be linked to their expression patterns, we analyzed published RNA-seq data sets, covering a broad range of samples: somatic or germ cells at different stages of developing male and female embryo (20 different conditions;(Guo et al. 2015)); pachytene spermatocytes and round spermatids from adult males (Lesch et al. 2016), and differentiated adult tissues (26 somatic tissues plus testis, which contains a fraction of germ cells;(Fagerberg et al. 2014). We first confirmed the negative relationship between intragenic recombination rate and gene expression level during meiosis (Fig 3A, S2A, S2D). We also confirmed that for both single cell data and bulk samples, the negative correlation between expression level and intragenic recombination rate is stronger in samples including germ cells than in somatic samples (Fig S3). This confirms that the intragenic crossover rate is affected specifically by expression in the germline.

**Fig 3:**
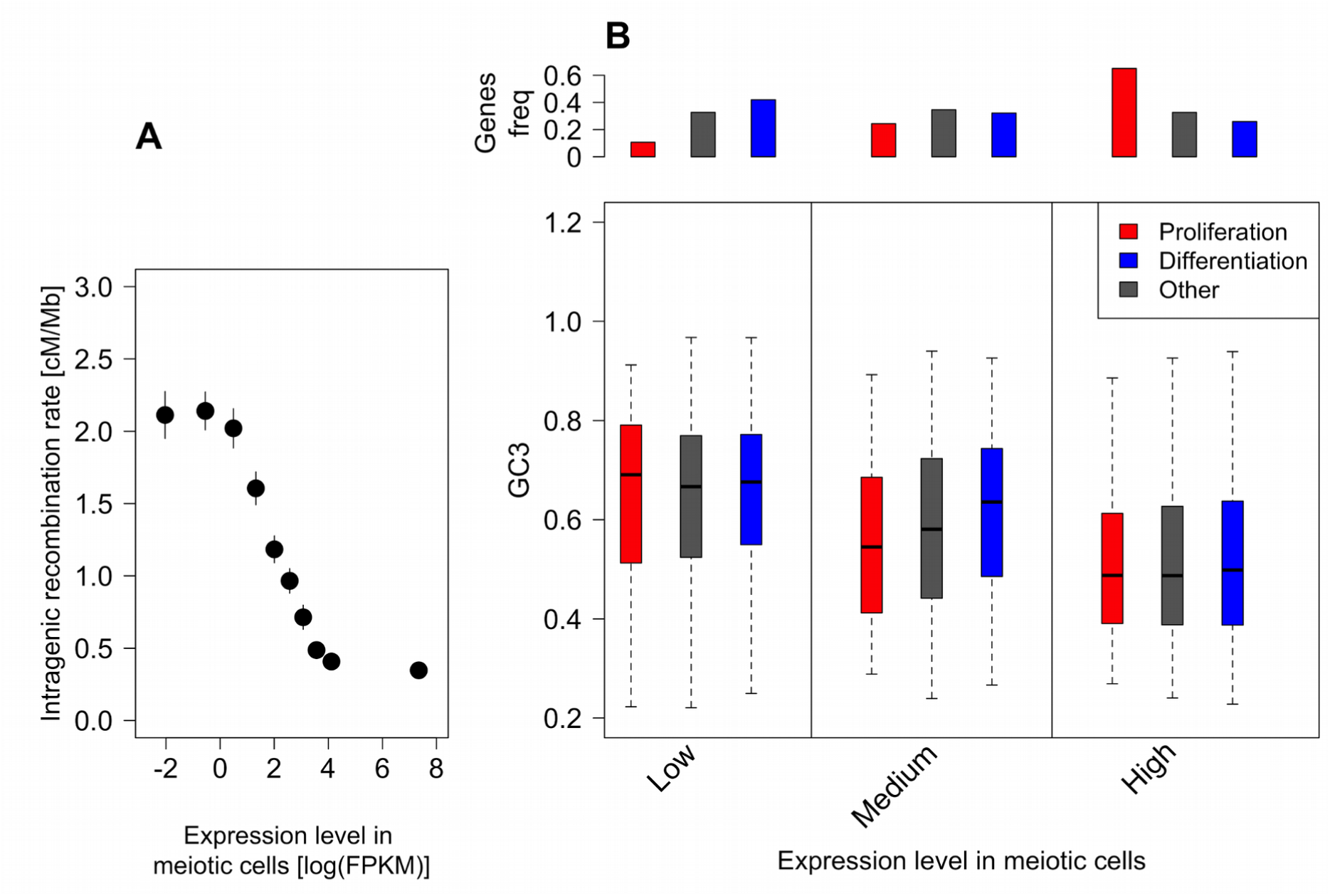
Variation in intragenic recombination rate and GC3 according to expression levels in meiotic cells. (A) Genes were classified according to their sex-averaged expression level in meiotic cells into 10 bins of equal sample size. The mean intragenic recombination rate was computed for each bin. Error bars represent the standard error of the mean. Similar results were obtained when analyzing separately expression levels in female or male meiotic cells (Fig S2A, S2D). (B) Variation in GC3 according to meiotic expression levels. Genes were first binned into 3 classes of equal sample size according to their sex-averaged expression level in meiotic cells (low: < 3.07 FPKM; high: >22.68 FPKM: medium: the others), and then split into three sets according to their functional category: “proliferation” (red), “differentiation” (blue), and “other” genes (grey). Boxplots display the distribution of GC3 for each functional category within each expression bin. Above barplots display the distribution of genes among expression bins for each functional category.

Many “proliferation” genes are involved in basic cellular functions, and hence, tend to be expressed at relatively high levels in many tissues and at all developmental stages. In particular, most of these genes are highly expressed in meiotic cells: 65% of “proliferation” genes are among the top 33% of genes with highest expression level (whereas only 11% are in the first tercile; Fig 3B). Conversely, only 26% of “differentiation” genes are highly expressed in meiotic cells, while 42% of are in the first tercile (Fig 3B).

This large proportion of “proliferation” genes with high meiotic expression levels can therefore explain why they tend to have relatively low intragenic recombination rate (Fig 2B), and hence, given the gBGC process, why they tend to have a lower GC3 (Fig 1C). To further test whether these differences in expression patterns could account for the difference in GC3 between “proliferation”, “differentiation” and “other genes”, we binned genes into three classes of increasing meiotic expression level. The distribution of GC3 is clearly shifted towards lower values for genes highly expressed at meiosis, compared to genes weakly expressed (average GC3 0.51 in the “high” category compared to 0.65 in the “low” category, p-value <2.10^*−16*^) (Fig 3B). However, there is no significant difference in the distribution of GC3 between “proliferation” and “differentiation” within bins of low or high expression (p-value = 0.68 and 0.15 respectively). Hence, the striking difference in synonymous codon usage between these functional categories (Figure 1C) disappears once the level of expression during meiosis is controlled for (Fig 3B).

Thus, differences in expression levels at meiosis may be responsible for differences in synonymous codon usage among gene categories in human, through the following causative chain: (i) The set of “proliferation” genes is enriched in genes highly expressed in meiosis. (ii) Because high expression at meiosis decreases the rate of crossovers, intragenic recombination rates are lower in the “proliferation” set. (iii) In turn, reduced intragenic recombination diminishes the effect of gBGC on exon base composition, and hence GC3 is lower in the set “proliferation” compared to “differentiation”.

To check whether this cascade of effects fully recapitulates the difference in synonymous codon usage between “proliferation” and “differentiation”, we investigated whether differences in SCU between functional categories is driven by expression level in cells undergoing meiosis, rather than by expression level in another cell type or tissue. We examined the relationship between GC3 and expression levels in a broad panel of cell and tissue conditions (Fig 4). As predicted by our model, expression levels in germ cells, either from single cell samples or from testis (which contains germ cells) are better predictors of GC3 than expression in all other somatic tissues. Strikingly, the levels of expression in primary germ cells is, on average, twice as informative than expression in somatic cells taken at comparable stage of development (Fig 4B). Among all individual samples, the strongest correlation between GC3 and expression level was found in male meiotic cells (pachytene spermatocytes, R^2^=6.3%, p-value<2.10^*−16*^). Female meiotic cells (primordial germ cells, PGC 17 W) showed a similar correlation level (R^2^=4.0%, p-value <2.10^*−16*^). As expected, the correlation is even stronger with sex-averaged meiotic expression level (R^2^=8.6%, p-value<2.10^*−16*^). Hence, these results confirm that the cell type for which gene expression level is the best predictor of GC3 (and therefore SCU) corresponds to meiotic cells.

**Fig 4:**
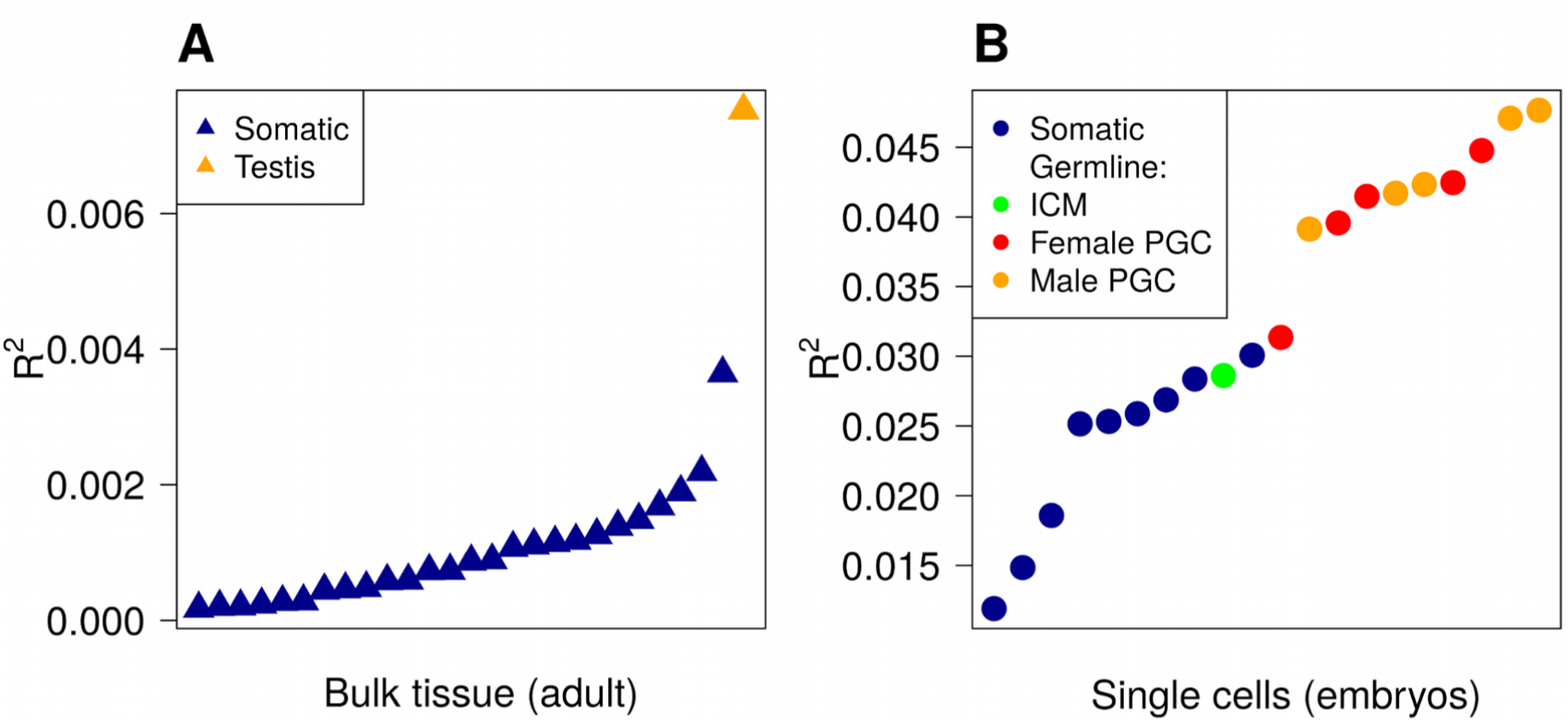
Correlation between expression level and GC3 in a panel of tissues and cell types. (A) bulk adult tissues data (Fagerberg et al. 2014) and (B) early embryo single cell data (Guo et al. 2015). These two subsets were obtained via very different protocols, which prevents direct cross-comparisons. Samples are sorted by increasing correlation coefficient (R^2^) between expression levels and GC3 (NB: all correlations are negative). Samples containing somatic cells are shown in blue; male germ cells in orange (testis or single cell) and female germ cells in red (PGC : primordial germ cells). The green point corresponds to cells from the inner cell mass (ICM) of the blastocysts, i.e. pluripotent cells from an early stage of development preceding the differentiation of germ cells.

### GC-content of non-coding regions and meiotic expression explain 70% of the variation in synonymous codon usage of human genes

Meiotic expression affects recombination rates along the entire gene (McVicker and Green 2010). Thus, the expression pattern is expected to affect gBGC intensity (and hence the GC-content) both in exons and in introns. Consistent with that prediction, the GC3 of human genes is strongly correlated to the GC-content of their introns (GCi, R^2^=62.7%, p-value <2.10*^−16^*). We build a linear model to quantify the relative contribution of the different parameters that covary with the GC3 of human genes (GCi, GC-flank, intragenic recombination rate, meiotic expression level, and “proliferation” or “differentiation” functional category). The analysis of variance demonstrates that GCi is by far the best predictor of GC3, but GC-flank, intragenic recombination rate and gene expression level during meiosis, also significantly improve the model (by 1%, 4% and 1.4% respectively, Table 1, ANOVA, p-values < 2.10*^−16^*). The integration of a categorical variable “differentiation” versus “proliferation” in the model significantly improves the model but its quantitative influence is minor (0.1%, p-value < 2.10*^−16^*, Table 1). Altogether, 68.2% of the variance in GC3 among human genes can be explained by the first four parameters (GCi, GC-flank, intragenic recombination rate, meiotic expression). Adding interaction terms to the linear model gives very similar results (70.4% variance explained, same levels of significance for all variables).

**Table 1:** Analysis of the variance of GC3 among individual genes. Variables included in the linear model are: GC-content of introns (GCi), GC-content of flanking regions (GC-flank), intragenic recombination rate (log scale), sex-averaged meiotic gene expression level (log scale) and functional category (“differentiation”, “proliferation” and “other”). Pairwise correlations (pairwise R^2^) were computed between GC3 and each of the other variables. Correlations of the model (model R^2^) were computed by adding variables sequentially.

| GC3 predictors | Pairwise $R^2$ | p-value | Model $R^2$ | F statistic | p-value |
| --- | --- | --- | --- | --- | --- |
| GCI | 62.7% | $<2.10^{-16}$ | 62.7% | 30232.4 | $<2.10^{-16}$ |
| + GC-flank | 48.1% | $<2.10^{-16}$ | 62.9% | 126.8 | $<2.10^{-16}$ |
| + Intragenic recombination rate | 12.8% | $<2.10^{-16}$ | 66.8% | 1453.3 | $<2.10^{-16}$ |
| + Expression level in meiosis | 8.3% | $<2.10^{-16}$ | 68.2% | 875.7 | $<2.10^{-16}$ |
| + Functional category | 1% | $<2.10^{-16}$ | 68.3% | 30.43 | $<2.10^{-16}$ |

## Discussion

In the human genome, gene sets that belong to different functional categories differ by their synonymous codon usage. Initially this pattern has been interpreted as evidence that the translation program was under tight control, notably to ensure a precise regulation of genes involved in cellular differentiation or proliferation (Gingold et al. 2014). According to this model, selection should optimize the match between the SCU of genes and tRNA abundances in the cells where they are expressed. However, the comparison of synonymous codon usage for amino-acids with single or multiple tRNA isoacceptors (Fig 1D) shows that the difference in SCU between functional categories does not result from constraints linked to tRNA abundance. In fact, variation in synonymous codon usage among functional categories is explained by one single dominant factor: the GC-content at third codon position (Fig 1B). The GC3 of human genes is strongly correlated to the GC-content of their introns and flanking regions (Table 1). This implies that variation in SCU results from a process that affects both coding and non-coding regions, and hence that it is not caused by translational selection.

Many lines of evidence indicate that large-scale variation in GC-content along chromosomes (isochores) is driven by the gBGC process, both in mammals and birds. First, there is direct evidence that recombination favors the transmission of GC-alleles over AT-alleles during meiosis (Odenthal-hesse et al. 2014; Arbeithuber et al. 2015; de Boer et al. 2015; Williams et al. 2015; Smeds et al. 2016). Second, the analysis of polymorphism and divergence at different physical scales (from kb to Mb) showed that recombination induces a fixation bias in favor of GC alleles (Duret and Arndt 2008; Clément and Arndt 2013; Munch et al. 2014; Pratto et al. 2014; Weber et al. 2014; Singhal et al. 2015; Glémin et al. 2015). Third, the gBGC model predicts that the GC-content of a given genomic segment should reflect its average long-term recombination rate over tens of million years (Duret & Arndt, 2008). Consistent with this prediction, analyses of ancestral genetic maps in the primate lineage revealed a very strong correlation between long-term recombination rates (in 1 Mb long windows) and stationary GC-content − R^2^=0.64; (Munch et al. 2014). The strong correlation between GC3 and GC-flank therefore implies that variation in synonymous codon usage is primarily driven by large-scale variation in long-term recombination rate.

Besides these regional fluctuations, recombination rates also vary at finer scale. In particular, recombination rates tend to be reduced within human genes compared to their flanking regions (Myers et al. 2005), and this decrease depends on the level of expression of genes during meiosis (McVicker and Green 2010) – see also Fig 3A. Hence, the gBGC model predicts that the GC3 of a gene should depend not only of the long-term recombination rate of the region where it is located, but also on its specific pattern of expression. And indeed, we observed that the difference in synonymous codon usage between “proliferation” and “differentiation” genes is not due to their preferential location in different classes of isochores, but to the fact that “proliferation” genes tend to be expressed a high level in meiotic cells, and therefore to have a reduced intragenic recombination rate (Fig 2, 3).

To test whether this observation holds true for other functional categories, we measured the average GC3, intragenic crossover rate and meiotic expression level of each GO gene set. As predicted by the gBGC model, we observed a very strong correlation between GC3 and the average intragenic recombination rate of GO gene sets (R^2^=0.51, Fig 5A). The variance in intragenic recombination rate, in turn, is very well explained by differences in meiotic expression levels among functional classes (R^2^=0.46, Fig 5B). As mentioned previously, these correlations measured on gene concatenates should be interpreted with caution because the different points are not independent (a same gene can belong to different GO categories). However, this analysis clearly shows that a large fraction of the variance in SCU observed among GO gene sets can be explained by variation in gBGC intensity, caused by variation in intragenic recombination rates, driven by differences expression patterns (Fig 5C). In agreement with the gBGC model, the intragenic recombination rate correlates with the base composition of the entire gene, including introns (Fig 5D). This observation clearly invalidates the hypothesis that the observed differences in SCU among functional categories might be driven by selection on codon usage.

**Fig 5:**
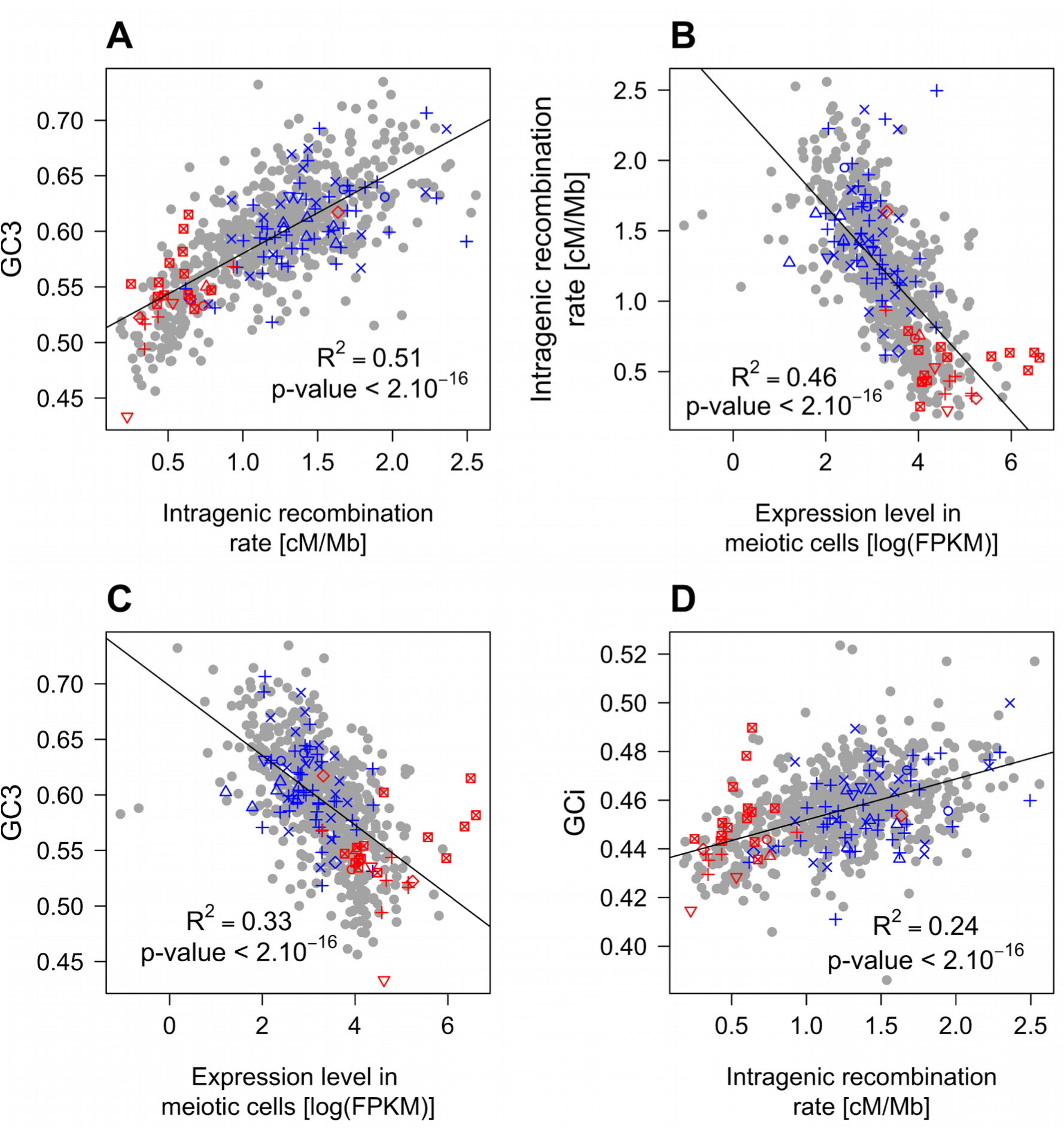
Relationships between GC-content, intragenic recombination rates and meiotic expression levels (sex-averaged) among functional gene categories. Average values of these parameters were computed for each GO gene set. We then measured correlations between these parameters: (A) Mean GC3 vs. mean intragenic recombination rate. (B) Mean intragenic recombination rate vs. mean expression level in meiotic cells. (C) Mean GC3 vs. mean expression level in meiotic cells. (D) Mean intronic GC-content (GCi) vs. mean intragenic recombination rate. GO gene sets associated to “proliferation” (red) or “differentiation” (blue) are displayed as in Fig 1. Similar results were obtained when analyzing separately expression levels in female or male meiosis (Fig S2).

The analysis of individual genes showed a much weaker correlation between GC3 and intragenic recombination rate (R^2^=12.8%; Table 1) than that observed with gene sets (R^2^=51%, Fig 5A). This difference can be explained by the fact that fine-scale recombination landscapes evolve very rapidly (Auton et al. 2012) and hence, present-day genetic maps are poor predictors of long-term intragenic recombination rate. For instance, although human and chimpanzee diverged only ∼7 million years ago, their recombination rates at the 10 kb scale are weakly correlated (R^2^=10%) (Auton et al. 2012). In gene set analyses, intragenic recombination rates are averaged over a large number of genes, which leads to reduce the variance caused by measurement errors and temporal fluctuations, and hence leads to increase the correlation with GC3 (Fig 5A). In absence of accurate estimates of long-term intragenic recombination rate of individual genes, we analyzed four indirect predictors: GCi, GC-flank, present-day intragenic recombination rate and meiotic expression levels. As expected, GCi is by far the best predictor of GC3 (Table 1). According to the gBGC model, if GCi was a perfect predictor of the long-term recombination rate within exons, then the other parameters should not appear as significant predictors of GC3. However, there is evidence that recombination rates differ between exons and introns (Kong et al. 2010). Moreover, whereas the base composition of exons is almost exclusively driven by base substitutions, introns are also affected by deletions and insertions (notably of transposable elements). Thus the base composition of introns does not perfectly reflect the long-term intensity of gBGC within exons. On the other hand, patterns of gene expression are well conserved among mammals (Brawand et al. 2011). Thus, expression levels measured in humans are expected to be good predictors of long-term average meiotic expression level, and thereby to provide some information on long-term intragenic recombination. This can explain why meiotic expression level appears as an important additional predictor of GC3 (Table 1). Altogether, these four variables explain 70% of the variance in GC3 of individual genes. In other words, the gBGC model can account for virtually all the variation in synonymous codon usage in the human genome.

It should be noted that co-variation between SCU and expression is generally considered as a typical signature of translational selection, and is often used to predict optimal codons (Duret 2002; Plotkin et al. 2004; Dos Reis and Wernisch 2009). However, as shown here, such correlations can also emerge as a result of a non-adaptive process. Given that gBGC is widespread in eukaryotes (Mancera et al. 2008; Capra and Pollard 2011; Pessia et al. 2012; de Boer et al. 2015; Williams et al. 2015; Smeds et al. 2016), it appears essential to take this process into account to interpret variation in synonymous codon usage (and more generally in base composition) among genes.

There is clear evidence the usage of synonymous codons is under selective pressure in some metazoan species (such as drosophila or nematode), which implies that it has a significant impact on the fitness of organisms – for review, see (Chamary et al., 2006; Duret, 2002; Plotkin & Kudla, 2011). It is *a priori* expected that codon usage should also affect translation efficiency (speed and accuracy) in mammals. However, our results show that selection on codon usage is not strong enough to counteract the impact of gBGC. In principle this does not exclude the hypothesis that the human genome might be subject to selection for translational efficiency: even if the GC-content of genes is driven by non-adaptive processes, there might be a selective pressure on the expression of tRNA genes to match the demand in synonymous codon usage. However, recent analyses of tRNA isoacceptors pools found no evidence for such variation (Schmitt et al. 2014; Rudolph et al. 2016). Moreover, we argue here that the peculiar base composition landscape induced by gBGC in the genomes of mammals and birds makes it impossible to match the tRNA pool to the demand in codon usage. Indeed, large-scale variation in recombination rates along the genome causes very strong variation in GC3 among genes, and this, whatever their functional category. In particular, “proliferative” genes, which are involved in basic cellular process, and are expressed at high levels in most tissues, show a very strong heterogeneity in GC3 (from 20% to almost 100%; Fig 1C). This implies that in any given cell, the set of highly expressed genes will show a very heterogeneous usage of synonymous codons. Hence, whatever the pool of tRNA available in that cell, there will be a large fraction of genes with a codon usage that does not match tRNA abundance. In other words, the heterogeneity of synonymous codon usage in mammalian genomes reflects a nonoptimal situation, caused the gBGC process, in which it is not possible to adapt the tRNA pool to the demand in codon usage of the transcriptome of any cell type.

### Material and Methods

#### Human protein coding genes

For each of the human protein coding genes in the Ensembl release 83 ((Yates et al. 2016); assembly GRCh38.p5), we identified a canonical transcript as defined in http://www.ensembl.org/Help/Glossary?id=346 (PERL script available in supplementary material on Zenodo platform: doi.org/10.5281/zenodo.229167). Mitochondrial genes were excluded from this analysis. Sequences of the remaining 19,766 canonical transcripts together with exons coordinates, were downloaded through the BioMart query interface (Smedley et al., 2015).

#### Recombination rates

Intragenic crossover rates were measured using the HapMap genetic map (The International HapMap Consortium, 2007). We chose this genetic map, which is based on the analysis of linkage disequilibrium in human populations, because its resolution (∼1 SNP per kb) is much higher than that of pedigree-based genetic maps (∼1 SNP per 10 kb) (Kong et al. 2010).

#### Definition of functional categories

The GO Term Accessions and GO domain were retrieved from Ensembl version 83 for the 19,766 genes. We retrieved biological process GO terms, counted the number of genes associated to each GO term and kept the ones that include at least 40 genes, except GO:0005515 that is too general to be informative (“protein binding” GO set, which includes 14,542 genes). This led to a final list of 687 GO gene sets. For each gene set, we concatenated coding sequences to compute the total codon usage, the relative synonymous codon usage (RSCU) and GC-content, and we also computed the average intragenic recombination rate and average expression levels (see below). The RSCU of a given codon corresponds to its frequency, normalized by its expected frequency if all corresponding synonymous codons were equally used (Sharp et al. 1986). For a given amino-acid (*x*), encoded by (*n_x_*) synonymous codons, the RSCU of its codon *y* is given by:

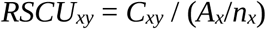

where C_xy_ is the number of occurrence of the codon *y* for amino-acid *x*, *A_x_* is the total number of occurrence of codons for the amino-acid *x*.

Following the classification used by (Gingold et al. 2014), we further defined two broad functional categories: “proliferation” and “differentiation”. GO terms containing the following keywords were associated to “proliferation”: “Chromatin modification”, “chromatin remodeling”, “mitotic cell cycle”, “mRNA metabolic process”, “negative regulation of cell cycle”, “nucleosome assembly”, “translation”. GO terms containing the following keywords were associated to “differentiation”: “Development”, “differentiation”, “cell adhesion”, “pattern specification”, “multicellular organism growth”, “angiogenesis”. Please note that GO terms corresponding to negative effects were excluded where appropriate (e.g. “negative regulation of proliferation” was not included in the “proliferation” category). Complete lists of GO terms are available at doi.org/10.5281/zenodo.229167

#### Analyses of individual genes

We also measured the codon usage of individual genes, to analyze covariations with their GC-content, expression levels and intragenic recombination rate. Owing to the low SNP density in human populations, the resolution of recombination maps is limited to about 5 kb (Myers et al, 2005). Because we investigate the relationship between GC3 and intragenic recombination rate, we selected genes that are long enough to measure recombination, i.e. at least 5 kb long (N=16,223 genes).

We defined three non-overlapping classes of genes according to their GO category: genes associated to at least one of the “proliferation” GO terms (N=1,008), genes associated to “differentiation” GO terms (N=2,833) and other genes (N=12,129). A group of 253 genes that were associated to both “proliferation” and “differentiation” GO terms were discarded from further analyses. The final dataset used in our analyses included 15,970 genes. In this dataset, there were 15,848 genes that contain at least one intron and for which we computed the GC content of intronic regions.

#### Expression data

Gene expression levels were collected from three publicly available human RNA-seq experiment datasets. The first one includes 27 differentiated adult tissues (Fagerberg et al. 2014; Kryuchkova-Mostacci and Robinson-Rechavi 2015) EBI accession number E-MTAB-1733). We downloaded normalized expression levels, already averaged across replicates, from Kryuchkova-Mostacci and Robinson-Rechavi (2015) (see supplementary information available on Zenodo platform at 10.5281/zenodo.229167). The second one is based on singlecell RNA-seq analysis, and includes 20 samples, corresponding to inner cell mass (ICM) of the blastocysts, and to primordial germ cells (PGC) and somatic cells, from male and female embryos at different development stages (4, 7 or 8, 10, 11 and 17 or 19 weeks, (Guo et al. 2015) GEO accession number GSE63818). We downloaded normalized expression levels from their dataset of pool-split PGCs (for more details see supplementary information at doi.org/10.5281/zenodo.229167). Female 17 weeks PGCs are entered in meiosis (Guo et al. 2015). This sample was therefore taken as representative of the transcriptome of meiotic cells in female. The third dataset corresponds to human male germ cells at pachytene spermatocytes (i.e. cells entering meiosis) and at round spermatids stages (post meiotic stage) ((Lesch et al. 2016); GEO accession number GSE68507, human RNA expression datasets GSM1673959, GSM1673963 GSM1673967, GSM1673971, GSM1673975 and GSM1673978). Guo and Lesch datasets include several replicates for each sample. We therefore computed the average expression levels over all replicates for each sample. The sex-averaged meiotic expression level was estimated by computing the mean of expression levels in female 17 weeks PGCs (Guo et al. 2015) and male spermatocytes or spermatids (Lesch et al, 2016). The correspondence between gene expression datasets and codon usage tables was based on Ensembl gene identifiers (Fagerberg and Lesch datasets), or on gene names (Guo dataset). In total, our analyses of expression levels were based on 15,305 genes (665 genes were absent from the Guo dataset).

#### Statistical analysis

Unless stated otherwise, reported R^2^ values correspond to Pearson correlation tests. R version 3.2.2 (R Core Team 2015) was used with Base package for statistical tests and graphics, plus ade4 library (Dray and Dufour 2007) for PCA analysis. The data and R scripts, which permit to reproduce the figures and tests presented here, are provided at: doi.org/10.5281/zenodo.229167

## Acknowledgement

This work was supported by French National Research Agency (ANR) grant DaSiRe (ANR-15-CE12-0010-01/DaSiRe). FP received a doctoral scholarship from Ecole Normale Supérieure de Lyon (http://www.ens-lyon.eu/). We thank Gaël Yvert for initiating the discussion and Adam Eyre-Walker for helpful suggestions on a first version of our manuscript.

